# The bi-directional extracellular electron transfer process aids iron cycling by *Geoalkalibacter halelectricus* in a highly saline-alkaline condition

**DOI:** 10.1101/2023.04.12.536630

**Authors:** Sukrampal Yadav, Chetan Sadhotra, Sunil A. Patil

## Abstract

Bi-directional extracellular electron transfer (EET) is crucial to upholding microbial metabolism with insoluble electron acceptors or donors in anoxic environments. Investigating bi-directional EET-capable microorganisms is desired to understand the cell-cell and microbe-mineral interactions and their role in mineral cycling besides leveraging their energy generation and conversion, biosensing, and bio-battery applications. Here, we report on iron cycling by haloalkaliphilic *Geoalkalibacter halelectricus* via bi-directional EET under haloalkaline conditions. It efficiently reduces Fe^3+^-oxide (Fe_2_O_3_) to Fe^0^ at a 2.29±0.07 mM/day rate linked to acetate oxidation via outward EET and oxidizes Fe^0^ to Fe^3+^ with a 0.038±0.002 mM/day rate via inward EET to reduce fumarate. Bioelectrochemical cultivation confirmed its outward and inward EET capabilities. It produced 895±23 μA/cm^2^ current by linking acetate oxidation to anode reduction via outward EET and reduced fumarate by drawing electrons from the cathode (−2.5±0.3 μA/cm^2^) via inward EET. The cyclic voltammograms of *G. halelectricu*s biofilms revealed redox moieties with different formal potentials, suggesting the involvement of different membrane components in bi-directional EET. The cyclic voltammetry and GC-MS analysis of the cell-free spent medium revealed the lack of soluble redox mediators, suggesting direct electron transfer by *G. halelecctricu*s in achieving bi-directional EET. By reporting on the first haloalkaliphilic bacterium capable of oxidizing and reducing insoluble Fe^0^ and Fe^3+^-oxide, respectively, this study advances the limited understanding of the metabolic capabilities of extremophiles to respire on insoluble electron acceptors or donors via bi-directional EET and invokes the possible role of *G. halelectricu*s in iron cycling in barely studied haloalkaline environments.

**Graphical abstract:** 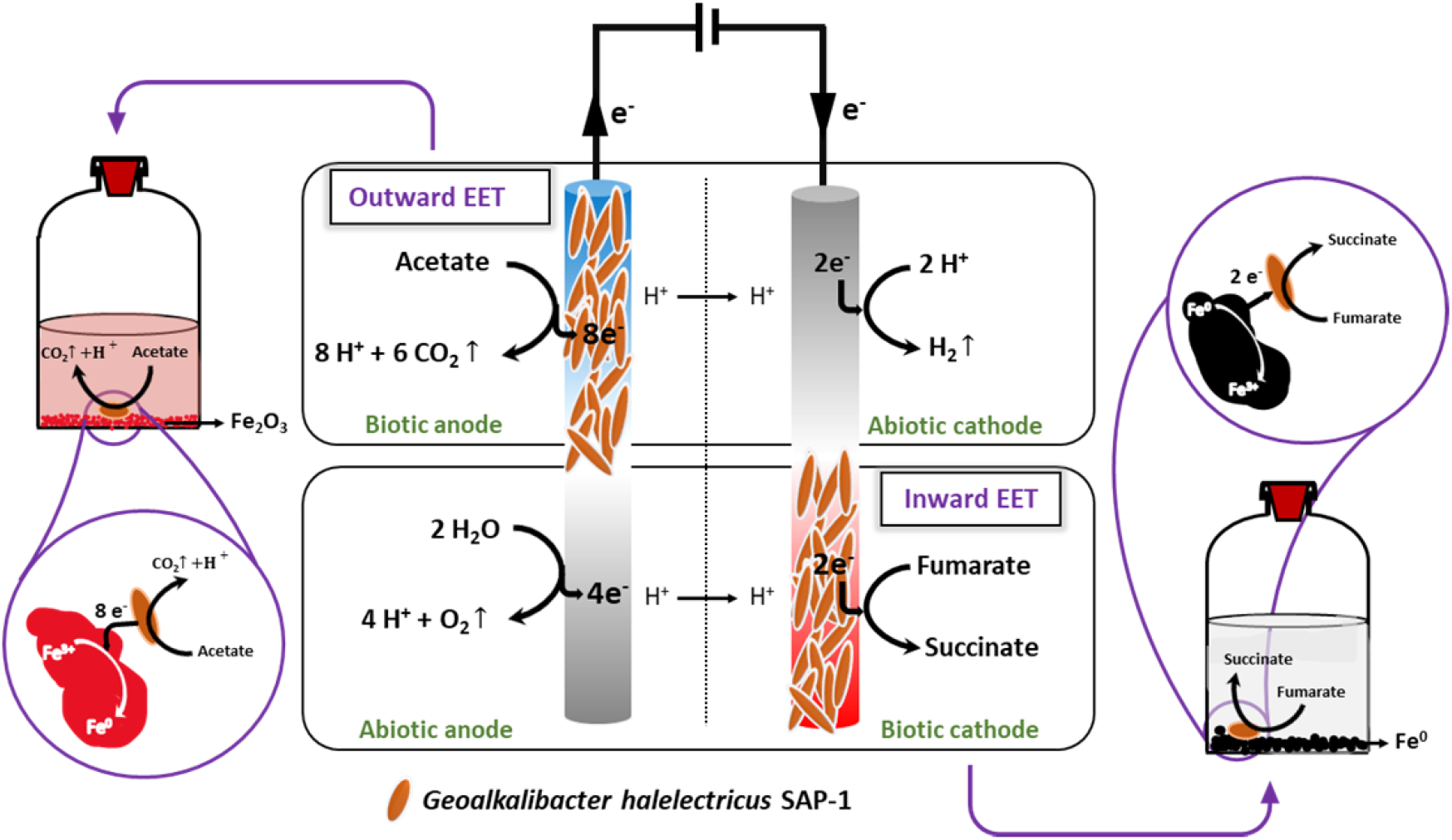

## 1. Introduction

Extracellular electron transfer (EET) aids microorganisms in achieving and maintaining their respiration on insoluble electron acceptors and donors in different anoxic environments (Patil et al., 2012; Lovley and Holmes, 2022). Using bioelectrochemical systems (BESs), EET between microorganisms and electrodes has been explored for developing various microbial electrochemical technologies (METs), which have lately emerged as a versatile platform for energy conversion and environmental remediation applications (Logan et al., 2019; Lovley and Holmes, 2022). The microorganisms capable of performing EET to exchange electrons with external conductive materials, such as metal oxides in natural environments and electrodes in the engineered BESs, are referred to as electroactive microorganisms (EAMs). The EET process is of two categories: outward, i.e., from the microbial cell or biofilm to the insoluble electron acceptor, and inward, i.e., from the insoluble electron donor to the microbial cell or biofilm (Nealson and Rowe, 2016; Reguera, 2018; Logan et al., 2019; Chaudhary et al., 2022). The outward EET is typically investigated by using the anode as the insoluble electron acceptor and the inward EET by providing the cathode as the insoluble electron donor in BESs (Holmes et al., 2022; Tremblay et al., 2017; Yu et al., 2018). For studying EET on natural insoluble electron donors or acceptors, oxidized metal compounds, such as Fe^3+^ oxide and Mn^4+^ oxide, and reduced metals or metal compounds, such as Fe^0^ and Mn^2+^ oxides, are employed (Dulay et al., 2020; Yee et al., 2020; Rotaru et al., 2021). Different microorganisms can also serve as electron donors or acceptors via direct interspecies electron transfer in natural anoxic environments and anaerobic digesters (Rotaru et al., 2014; Cheng and Call, 2016; Baek et al., 2018; Rotaru et al., 2019; Holmes et al., 2022). Though several microorganisms have been reported for either outward or inward EET (Kiran and Patil, 2019; Logan et al., 2019), few are known for their bi-directional EET, i.e., both outward and inward EET, capabilities. Microorganisms with bi-directional EET capabilities not only play important roles in the minerals or element cycling, microbially induced corrosion, and interspecies electron transfer processes in anoxic environments but are also useful in developing biosensing, energy storage alternatives or bio-battery, and bioelectronic applications (Jiang and Zeng, 2019; Li et al., 2021). Studying microbial diversity capable of bi-directional EET is thus essential to advance the understanding of this environmentally relevant and applied microbial phenomenon.

A few studies reported on the bi-directional EET of mixed and pure microbial cultures. In most outward EET demonstration studies, acetate has been used as the electron donor source, whereas different types of electron acceptors have been used for demonstrating the inward EET process in BESs. For instance, the model EAM *Geobacter sulfurreducens* is well-studied for outward EET with the anode electrode and for inward EET process from the cathode to reduce protons, fumarate, and soluble U(VI) to hydrogen, succinate, and U(IV), respectively (Gregory et al., 2004; Gregory and Lovley, 2005; Soussan et al., 2013). *Shewanella oneidensis*, another model EAM, also exhibit bi-directional EET to grow by using an anode as the electron acceptor (Xu et al., 2018) and to reduce nitrobenzene to aniline with Fe^2+^ (Luan et al., 2015) as the sole electron donor via flavin-based bi-directional EET (Okamoto et al., 2014; Zou et al., 2019). Other microbes reported for mediated bi-directional EET include fermentative facultative anaerobes *Klebsiella pneumoniae* and *Escherichia coli* (Zhang et al., 2006; Zhang et al., 2008; Harrington et al., 2015). *Alcaligenes faecalis*, a facultative anaerobic bacterium, performs outward and inward EET to catalyse autotrophic denitrification and dissimilatory nitrate reduction to ammonium reactions (Yu et al., 2018). The mixed microbial electroactive biofilms dominated by diverse groups have also been reported for bi-directional EET to attain charge-discharge behaviour or energy-generating and energy-storing reactions (Izadi et al., 2021; Li et al., 2021; Mickol et al., 2021), and bioelectrochemical denitrification (Liang et al., 2019; Pous et al., 2016).

Notably, the direct electron transfer-based bi-directional EET is reported only for *Geobacter* sp., *Shewanella* sp. (Xie et al., 2021), and *Desulfurivibrio alkaliphilus* (Izadi and Schröder, 2022). Moreover, most bi-directional EET studies have been conducted in BESs with non-extreme microorganisms and with the electrode serving as either an electron donor or acceptor source and barely with natural insoluble electron donors and acceptors, such as Fe and Mn elemental and oxide forms (Jiang and Zeng, 2019; Xie et al., 2021). Thus, understanding of the role of bi-directional EET-capable microorganisms, particularly extreme microbes, in cycling elements such as Fe in anoxic conditions remains poor. To this end, the present work pertains to studying iron cycling by a recently reported haloalkaliphilic *Geoalkalibacter halelectricus* SAP-1 via bi-directional EET under highly saline-alkaline (pH 9.5 and salinity 20 g/L) conditions. It is an anaerobic, non-spore-forming, rod-shaped, Gram-negative, and heterotrophic EET-capable bacterium isolated from the electroactive biofilm enriched from haloalkaline lake sediments (Yadav et al., 2022). The bi-directional EET capabilities were checked with Fe^3+^-oxide and Fe^0^ as the sole insoluble electron acceptor and donor under strictly anoxic conditions in biochemical experiments. The outward and inward EET capability of *G. halelectricus* was further confirmed through an electrochemical approach involving cultivation with an anode and cathode as the sole electron acceptor and donor. Next, cyclic voltammetry was applied to check the EET mode used by *G. halelectricus* in the bi-directional EET process. In addition, the microbial attachment over the electrode and iron particle surfaces was observed with SEM, and EDS was used for energy-dispersion-based element identification after biochemical and electrochemical cultivation experiments.

## 2. Experimental procedures

### 2.1 Microbial culture and growth conditions

*Geoalkalibacter halelectricus* SAP-1 (=JCM 35356, =MTCC 13188) culture maintained in the laboratory was used in this study. All experiments, including media preparation and incubations, were conducted under aseptic and anoxic conditions and at a controlled temperature of 30°C, if not stated otherwise. A modified M9 medium with pH 9.5 and salinity 20 gNaCl/L and containing (per L of distilled water) 4.33 g Na_2_HPO_4_, 2.69 g NaH_2_PO_4_, 20 g NaCl, 4.3 g Na_2_CO_3_, 0.13 g KCl, 0.31 g NH_4_Cl, 12.5 ml vitamins, 12.5 ml trace elements, and suitable electron donor or acceptor according to the tested experimental conditions, was used for culturing and maintaining *G. halelectricus* (Yadav and Patil, 2020; Yadav et al., 2022). All chemicals and reagents were of analytical or molecular grade and obtained from Merck and Sigma-Aldrich.

### 2.2 Microbial activity with Fe^3+^ oxide and Fe^0^ as the electron acceptor and donor

These experiments were conducted in triplicates in serum flasks (100 ml capacity with 60 ml working volume) under strict anoxic and haloalkaline growth medium conditions. Fe^3+^ oxide and elemental iron (Fe^0^) were used as the sole insoluble electron acceptor and donor sources to conduct outward and inward EET experiments, respectively. For the outward EET experiment, a modified M9 medium containing acetate (10 mM) and Fe_2_O_3_ (hematite; 30 mM) as electron donor and acceptor, respectively, was used. Microbial inoculum (10 % of working volume) with an optical density of ∼1.0 was added to the serum flasks. The bacterial cells were collected via centrifugation on complete acetate consumption in outward EET experiments. It was followed by pellet washing with modified M9 medium buffer thrice. Then, the washed bacterial cells were used as the inoculum source in inward EET experiments. These experiments were conducted only to monitor the microbial activity for Fe^0^ oxidation; hence, no additional carbon or electron source was supplemented in the medium. The medium contained solid iron powder (Fe^0^; ∼224 mM) as the sole electron donor and fumarate (30 mM) as the terminal electron acceptor. The abiotic control experiments were conducted to check the abiotic Fe^3+^ reduction and Fe^0^ oxidation in the corresponding experimental conditions. Another set of control experiments contained Fe^3+^ and microbial inoculum but lacked acetate and was conducted to check the possibility of Fe^3+^ reduction linked to the oxidation of any components other than acetate, specifically ammonium ions in the medium. The concentration of all electron donors and acceptors, viz. acetate and Fe^3+^ ions for outward EET and Fe^3+^ ions and fumarate for inward EET, were monitored on alternate days.

In addition, cyclic voltammetry (CV) and Gas Chromatography-Mass Spectrophotometric (GC-MS) analysis of the spent medium was conducted to check the presence of any soluble redox mediator that can aid the EET process. CV was conducted with the filtered cell-free spent medium of outward and inward EET experiments. The GC-MS analysis was conducted for the growth medium and filtered cell-free spent medium of both outward and inward EET experiments. For GC-MS, dichloromethane and methanol with 1:1 ratio was used to extract the polar fraction. Then, the concentrated solution (∼0.5 ml) was placed in GC vials, and 1 μl of that was injected into a gas chromatograph coupled to the mass detector (GC-MS, 7890B GC, 5977C MSD, Agilent Technologies). It consisted HP-5MS capillary column to separate the analytes and used helium as carrier gas with a flow rate of 1 ml/min. The GC oven was programmed between 40 to 320 °C at a ramp rate of 5 °C/min (Kumar et al., 2021).

### 2.3 Bi-directional EET confirmation with electrodes as the solid-state electron acceptor and donor

These experiments were conducted in three independent bioelectrochemical reactor replicates (R1, R2 and R3) by following the methodology explained below.

#### 2.2.1 Electrochemical setup and microbial biofilm growth at the electrode surface

The electrochemical cultivation approach was used to grow the *G. halelectricus* biofilm at the anode surface, which served as the sole terminal electron acceptor (Singh et al., 2022). For this, potentiostatically controlled two-chambered reactors with a three-electrode configuration were used. Briefly, two electrode chambers were separated by a proton exchange membrane (Nafion, Sigma-Aldrich), and each chamber had a working volume of 200 ml. Graphite plates with a projected surface area of 7.0125 cm^2^ were used as anode and cathode electrodes. The membrane and graphite electrodes were pre-treated before use according to protocols mentioned elsewhere (Singh et al., 2022). An Ag/AgCl (3.5 M KCl,0.205 V vs. SHE, RE-1B, Biologic Science Instruments, France) reference electrode was placed in the anode chamber to monitor the electrode potentials. A modified M9 medium with 20 g/L salinity, 9.5 pH and acetate (20 mM) as a sole carbon and electron donor source was used as an electrolyte or growth medium in the anode chamber. It lacked any other soluble or insoluble electron acceptor except the solid-state electrode. The catholyte contained only buffer components of the same medium. The electrochemical cultivation experiments were conducted by polarizing the working electrode at 0.2 V vs. Ag/AgCl, and the bioelectrocatalytic current response at a fixed time interval of 2 min was recorded using the chronoamperometry (CA) technique. The polarized anode thus acted as an analogue to Fe^3+^-oxide and created a considerable potential difference between the electron donor, i.e., acetate, and the electron acceptor, i.e., anode. The microbial inoculum (OD_600_ ∼1.0) size was 10 % of the working volume of the anode chamber. To develop a well-established biofilm at the anode surface (i.e., bioanode), at least three batch growth cycles were conducted by replenishing the entire spent medium with a fresh medium at the end of each batch cycle. Abiotic and biotic electrochemical control experiments were also conducted. The abiotic control lacked any microorganisms and aimed to examine the possibility of electrochemical current generation through abiotic acetate oxidation. Whereas the biotic control contained microbial inoculum but lacked acetate and aimed to examine the possibility of bioelectrocatalytic current generation through oxidation of components other than acetate, specifically ammonium ions, in the medium.

To investigate the inward EET of *G. halelectricus*, the previously developed bioanode was switched to a biocathode (i.e., microbial biofilm at the cathode surface) operation mode to check its capability to uptake or draw electrons directly from the cathode and use them for the fumarate reduction reaction. After 3-4 batch cycles, the growth medium was replenished by an acetate-free modified M9 medium (pH 9.5 and salinity 20 g/L) containing 30 mM fumarate as a sole terminal electron acceptor. No electron source other than the polarized cathode was used. To favour the fumarate reduction reaction under these conditions, the applied cathode potential was set to −0.650 V (vs. Ag/AgCl), and bioelectrocatalytic reduction current was recorded at a fixed time interval of 2 min using CA. The applied cathodic potential was chosen in a manner that enables fumarate reduction and creates a sufficient potential difference between the electron donor, i.e., polarized cathode and electron acceptor, i.e., fumarate (−0.025 V vs. Ag/AgCl at pH 9.5). The anode chamber was operated under oxic conditions to favour the water oxidation reaction and provide electrons to the cathode. For this purpose, a mixed metal oxide-coated titanium plate was used as the anode. An abiotic electrochemical control experiment with a bare cathode (i.e., lacking cathodic microbial biofilm) was conducted to check the possibility of electrochemical fumarate reduction. The microbial growth medium was purged with 99.999% N_2_ gas (Sigma Gases, India) for at least 20 min before starting any experiment. The pH and the concentrations of acetate and fumarate in the bulk phase and hydrogen in the headspace were monitored at regular intervals.

#### 2.2.2 Electrochemical analysis of the outward and inward EET by G. halelectricus biofilms

To understand the EET mode and formal potentials of the redox-active moieties used by *G. halelectricus* in both outward and inward EET, CV analysis at different experimental conditions was conducted. For outward EET, CVs were recorded before and immediately after microbial inoculation, during substrate turnover and non-turnover conditions, and with the fresh electrode in the filtered cell-free spent medium by ramping the working electrode potential in the range from −0.4 to +0.6 V at 1 mV/s scan rate. For inward EET, CVs were recorded before and immediately after microbial inoculation, at substrate turnover condition, and with the filtered spent medium by ramping the working electrode potential in the range from −1.0 to 0.0 V at 1 mV/s scan rate. The changes in oxidative and reductive faradic current were recorded at a particular ramped potential. The first derivative analysis was applied to find the mid-point or formal potential of the redox-active moieties or peaks in the recorded CVs.

### 2.4 Characterization and analysis

The microbial cell attachment over the Fe^3+^ oxide and Fe^0^ surface and biofilm growth on the electrode surfaces were visualized using scanning electron microscopy (SEM), as per the procedure explained elsewhere (Yadav and Patil, 2020). The elemental analysis using energy-dispersive X-ray spectroscopy (EDS) was conducted to confirm the Fe particle formation observed during SEM analysis. For SEM and EDS analysis of the bulk phase, the liquid samples were passed through isopore membrane filters (Sigma-Aldrich), and the filters were subjected to observations. The acetate concentration was analysed by HPLC (Agilent 1260 Infinity II, RID Detector, Hiplex H column, 5 μM H_2_SO_4_ as mobile phase, flow rate 0.5 ml/min, Temperature 50°C). H_2_ gas production in cathodic experiments was monitored using GC-TCD (Agilent 490 MicroGC). Spectrophotometric detection methods, namely, phenanthroline, phenate, hydrochloric acid and Griess reagent, were used to quantify Fe^3+^, ammonium, nitrate, and nitrite concentrations, respectively. For quantifying the fumarate concentration, a fumarate assay kit (MAK060, Sigma-Aldrich) was used as per the manufacturer’s guide.

## 3. Results and Discussion

### 3.1 *G. halelectricus* is capable of utilizing insoluble Fe^3+^-oxide and Fe^0^ as the sole electron acceptor and donor

*G. halelectricus* grew well with acetate and Fe^3+^ oxide as the sole electron donor and acceptor, respectively. Acetate oxidation and Fe^3+^ reduction occurred simultaneously (**Figure 1a**). *G. halelectricus* oxidized acetate efficiently (i.e., from initial 10 mM to final 1.17±0.14 mM) to reduce Fe^3+^ from initial 30 mM to final 4.23±0.47 mM (86.14±0.6 %) within 12 days of the incubation period. It reduced Fe^3+^ at a 2.29 ± 0.07 mM/day rate. The microbial Fe^3+^ reduction occurred via Fe^2+^ as an intermediate, which was observed transiently and further reduced to Fe^0^. No decrease in Fe^3+^ was observed in abiotic control experiments (**Figure 1a**). In addition, Fe^3+^ reduction was not observed in another control experiment lacking acetate (**Figure S1**), suggesting the inability of *G. halelectricus* to use any other component from the medium as the electron source to reduce Fe^3+^. It confirmed the microbial role in Fe^3+^ reduction coupled with acetate oxidation in main biochemical experiments. The ability to grow using insoluble Fe^3+^ as the sole electron acceptor suggests the outward EET capabilities of *G. halelectricus* to achieve its respiration. Quantifying the exact Fe^2+^ concentration was not possible because of its transient production and further rapid reduction to Fe^0^. So, only the substrate (Fe^3+^) concentration was quantified throughout the experiment.

**Figure 1:**
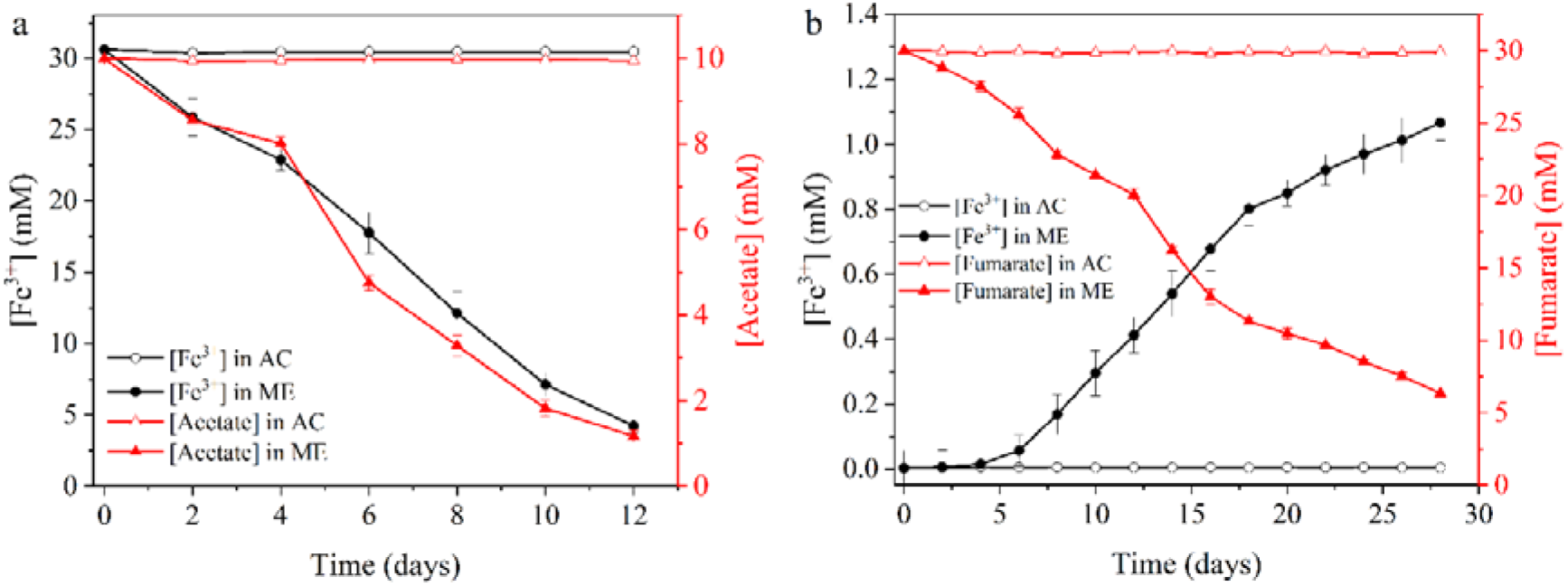
Cultivation of *Geoalkalibacter halelectricus* on Fe^3+^-oxide (hematite) as the sole electron acceptor (electron donor: acetate) (a) and Fe^0^ as the sole electron donor (electron acceptor: fumarate) (b) (n=3). ME – Main experiments inoculated with microorganisms; AC– Abiotic Control.

*G. halelectricus* also showed Fe^0^ oxidation activity (measured as Fe^3+^, the final Fe^0^ oxidation product) linked to fumarate reduction (**Figure 1b**). Over an incubation period of 28 days, it oxidized limited Fe^0^ resulting in 1.07±0.05 mM Fe^3+^ concentration at 0.038±0.001 mM/day rate. The fumarate concentration decreased from 30 mM to 6.34±0.21 mM during the same incubation period. These coupled redox reactions for Fe^0^ oxidation, only in the case of inoculated sets but not in abiotic control (**Figure 1b**), suggest the inward EET capabilities of *G. halelectricus* to maintain its metabolic activity.

The digital photographs of the flasks captured at the start and the end of outward experiments revealed a colour change in the growth medium. The medium ultimately turned colour-less (with iron particles settled at the bottom) (**Figure 2d**) from initial yellowish (due to Fe^3+^) in the Fe^3+^-oxide reduction experiments (**Figure 2a**). The SEM analysis of the samples from the outward EET experiments revealed the attachment of *G. halelectricus* cells to the Fe^3+^ oxide particles (**Figure 2e**). In contrast, bare Fe^3+^ oxide particles without microbial growth or cell attachment were observed in the case of abiotic control (**Figure 2b**). In the EDS analysis of *G. halelectricus* grown with Fe^3+^ oxide, >48.4 % of iron besides 51.2 % carbon was detected (**Figure 2f, Table S1**), thereby confirming the formation of Fe^0^ particles. Similar data were obtained for the samples of other replicates (**Figure S2, Table S1**). No Fe^0^ particles but only carbon (C, 100% atomic composition) were observed in the abiotic control (**Figure 2c, Table S1**). The carbon peaks originated from the carbonaceous isopore membrane filters used for the EDS sample preparation in all cases (**Figures 2c, 2f, S2, and Table S1**). The negligible Fe^3+^ and Fe^2+^ concentrations and EDS analysis of the particles after the completion of experiments suggested the occurrence of mainly elemental iron particles. Similar to the outward EET, the colour of the growth medium became yellowish (due to the production of Fe^3+^) (**Figure 3b**) from colour-less (with iron particles settled at the bottom) (**Figure 3a**) in Fe^0^ oxidation experiments. The SEM analysis of the samples from the inward EET experiments revealed the attachment of *G. halelectricus* cells to the Fe^0^ particle surfaces (**Figures 3c and d**). No redox peak or activity was observed in the CVs recorded in the filtered cell-free spent media of outward and inward EET experiments (**Figure S3**). In addition, the mass spectrographs of the growth medium and spent media of outward and inward EET experiments revealed the absence of any commonly used redox mediators, such as flavins (700 m/z), riboflavin (242 m/z), and phenazine (180 m/z) (**Figure S4)**. Thus, both CV and GC-MS analyses suggested the absence of any soluble redox mediator secreted by *G. halelectricus* to achieve Fe^3+^ oxide reduction and Fe^0^ oxidation processes.

**Figure 2:**
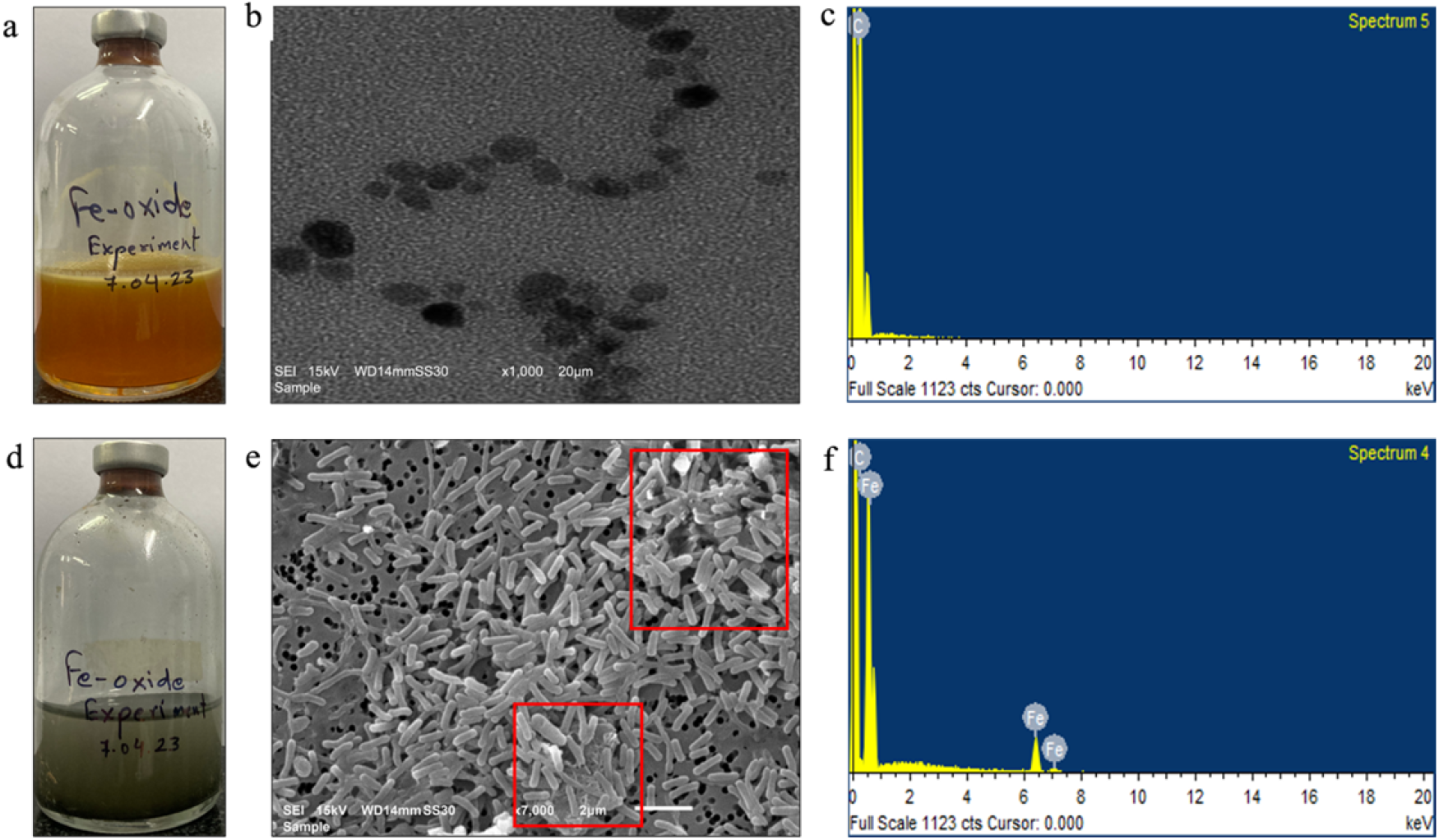
Digital photograph of serum flask (a), SEM image (b) and EDS elemental profile (c) of abiotic control compared to the digital photograh of serum flask (d), SEM image (e) and EDS elemental profile (f) of *G. halelectricus* cultivated on Fe^3+^ oxide in outward EET experiments. The initial yellowish growth medium color turned colourless at the end of experiment (d). Fe^3+^ oxide particles are marked in the red box. The carbon peaks observed in the elemental profile originate from the carbonaceous filter used in sample preparation for EDS analysis.

**Figure 3:**
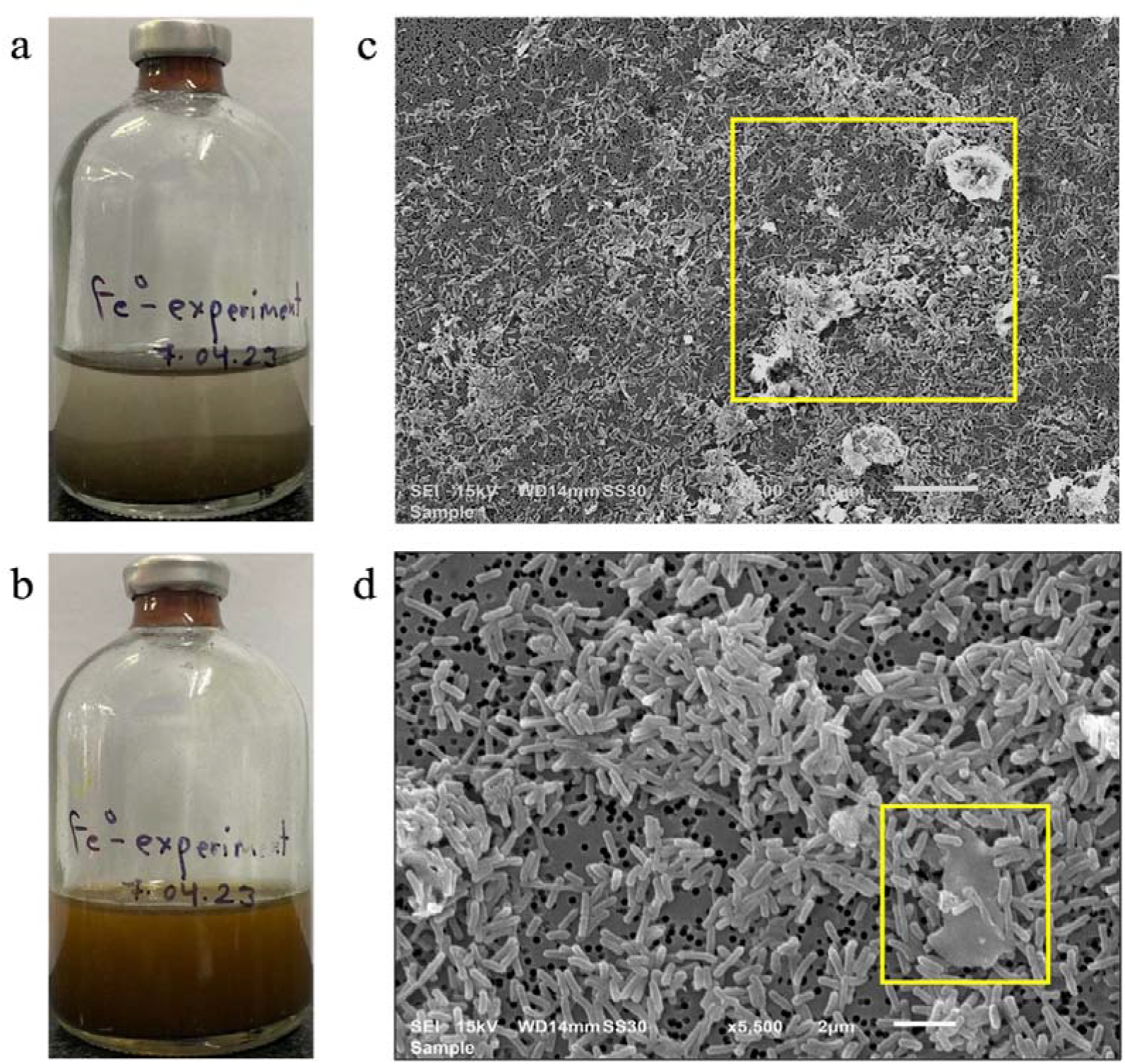
Digital photographs of representative serum flasks showing the change in medium colour at the start (a) and end (b) of the experiment, corresponding to Fe^0^ oxidation to Fe^3+^. SEM images showing attachment of *G. halelectricus* cells to Fe^0^ surfaces (at 10 μm (a) and 2 μm (b) scales) in inward EET experiments with Fe^0^ as the sole electron donor. Fe^0^ particles are marked in the yellow boxes.

### 3.2 Electrochemical cultivation of *G. halelectricus* with electrodes as the electron acceptor and donor confirms its bi-directional EET capability

#### 3.2.1 Outward EET capability

The electrochemical cultivation of *G. halelectricus* at an applied potential of 0.2 V led to the growth and development of a brownish-coloured biofilm at the anode electrode. It produced bioelectrocatalytic current by linking acetate oxidation to anode reduction (**Figure 4a, Figures S5a and S5b for replicate reactors**). It reached a maximum current density of 895±23 μA/cm^2^, which subsequently decreased along with the decrease in acetate concentration (to about 2.3±0.11 mM) in the batch experiments. The current production resumed instantaneously on replenishment of the spent medium with fresh medium containing acetate in each batch cycle. Neither acetate consumption nor substrate oxidation current generation was observed in the abiotic control experiment (**Figure 4b**). The absence of any faradic current production in abiotic control confirms the bioelectrocatalytic current generation capability of *G. halelectricus* by oxidising acetate and respiring on the anode as the sole electron acceptor via outward EET. Similarly, the absence of any faradic current production in the biotic control lacking acetate confirms the incapabilities of *G. halelectricus* to respire on the anode using any other component, especially ammonium as the electron source in the medium (**Figure 4b**).

**Figure 4:**
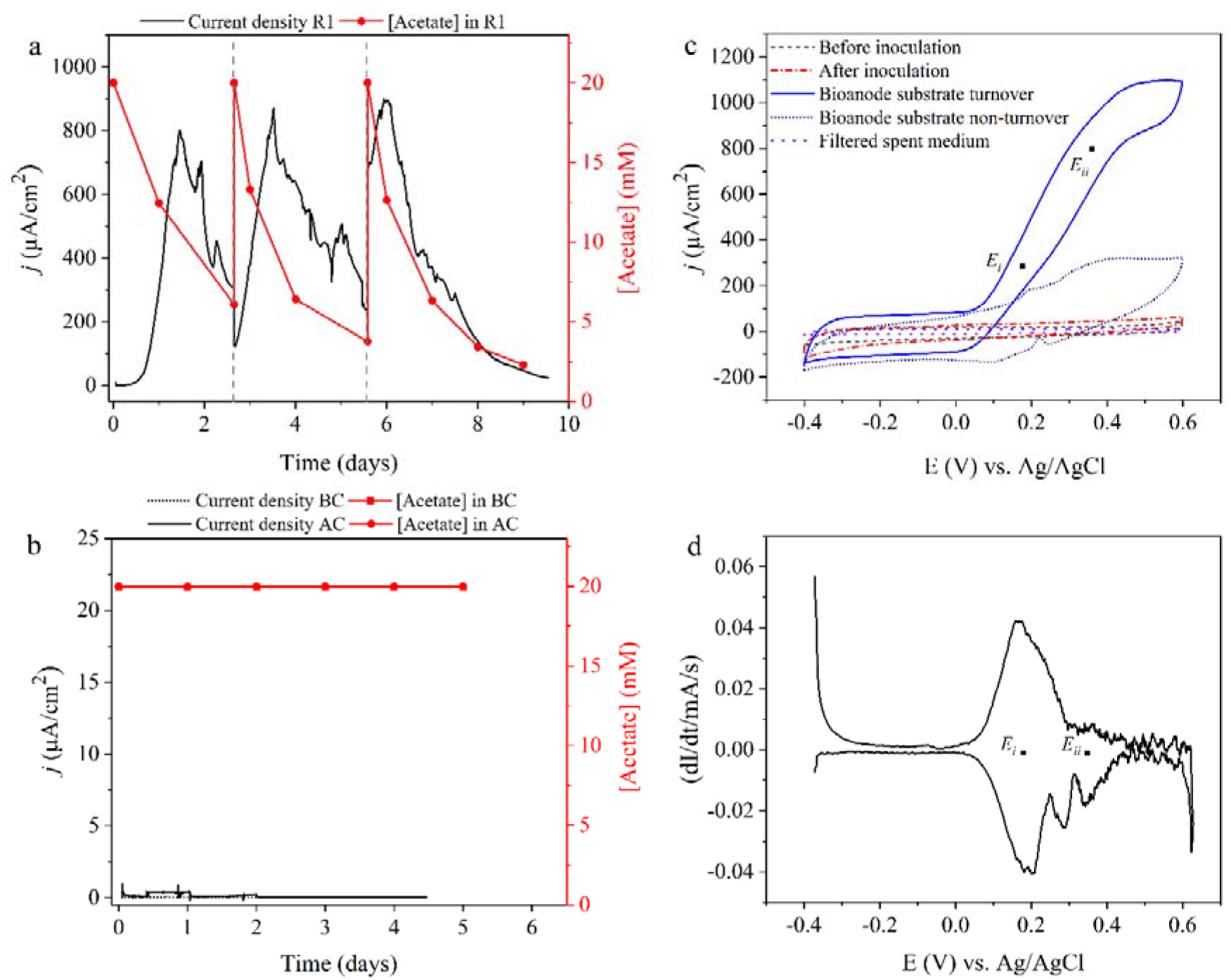
Electrochemical investigation of outward EET capability of *Geoalkalibacter halelectricus* with anode as the sole electron acceptor (substrate/electron donor: acetate). Bioelectrocatalytic anodic current generation and acetate concentrations profiles of representative bioelectrochemical reactor R1 (a) and control experiments (b), cyclic voltammograms (CVs) at different conditions viz, before and after inoculation, during substrate turnover and non-turnover, and filtered cell-free spent medium recorded for the same reactor (c), and the first derivative of CV recorded at substrate turnover condition (d). AC – Abiotic Control, BC – Biotic Control.

The cyclic voltammograms (CVs) of the bioanode recorded at substrate turnover and non-turnover conditions showed signatory curves with a prominent redox activity at a mid-point or formal potential of ∼0.181 V vs. Ag/AgCl (**Figure 4c and 4d; Figure S6**). Another set of redox peaks with a mid-point potential of ∼0.349 V was also observed. No redox activity was observed in the anodic CVs recorded before and immediately after the inoculation suggesting the absence of any redox-active compound or moiety in the modified M9 medium and over the abiotic electrode surface. In addition, no redox activity was observed in the CVs recorded with filtered cell-free spent medium, suggesting that microbes did not produce any soluble redox mediators. These results suggest that *G. halelectricus* uses most likely a direct mode of EET to achieve respiration with the anode. It is achieved via the outward EET mechanism by expressing and arranging the electron transport chain components according to ascending reduction potential order (Patil et al., 2012; Shi et al., 2016). The mid-point potential of the redox moiety (0.181 V) observed in bioanode CVs is slightly lower than the applied electrode potential (0.2 V) and much higher than the acetate oxidation potential (−0.344 V) under here-tested experimental haloalkaline conditions. The large potential difference between the electron donor and acceptor favours the electron array from the microbial cell membrane to the anode and establishes the outward EET process.

#### 3.2.2 Inward EET capability

On switching the polarity to the cathodic mode by applying −0.65 V vs. Ag/AgCl and without any electron acceptor (fumarate), a stable reduction current almost close to zero was observed for six days. With the addition of fumarate, the reduction current increased (−2.5±0.3 μA/cm^2^), which was confirmed by operating reactors for three batch cycles (**Figure 5a, Figures S7a and S7b for replicate reactors**). The reduction current was correlated with the decrease in fumarate concentration (6.85±0.83 mM) with its reduction efficiency of 74.8±2.17 %. Neither decrease in fumarate concentration nor electrocatalytic current withdrawal from the cathode was observed in the abiotic control experiments (**Figure 5b**). In addition, H_2_ was not detected in the cathodic chamber headspace of the abiotic control and the main biocathode experiments at the applied cathode potential. The CVs recorded at different conditions: before and immediately after inoculation and turnover condition with electron acceptor (i.e., fumarate), also suggest the inward EET capabilities in *G. halelectricus*. Two redox moieties with the formal potential of −0.450 and −0.580 V vs. Ag/AgCl were observed in biocathode CVs (**Figure 5c and 5d; Figure S8**). In comparison, no redox activity or peak was observed in the CVs recorded in the filtered cell-free spent medium, suggesting a lack of any microbially-secreted soluble redox mediator. In addition, the CVs of the abiotic cathode lacked typical hydrogen electrocatalysis traces at lower potentials. These CV observations, besides fumarate reduction linked to the cathodic reduction current in CA experiments, suggest most likely the direct mode of electron uptake rather than indirect EET from the cathode by *G. halelectricus*. In this case, *G. halelectricus* might have expressed the membrane components with formal potentials of −0.450 and −0.550 V in drawing electrons from the cathode poised at −0.650 V and reducing fumarate (−0.025 V at 9.5 pH), with descending reduction potential order (Jiang and Zeng, 2019; Xie et al., 2021). The redox moieties with a formal potential close to −0.450 and −0.550 V have been reported earlier for fumarate-reducing *Geobacter sulfurreducens* in inward EET experiments (Pous et al., 2016; Liang et al., 2019; Li et al., 2021).

**Figure 5:**
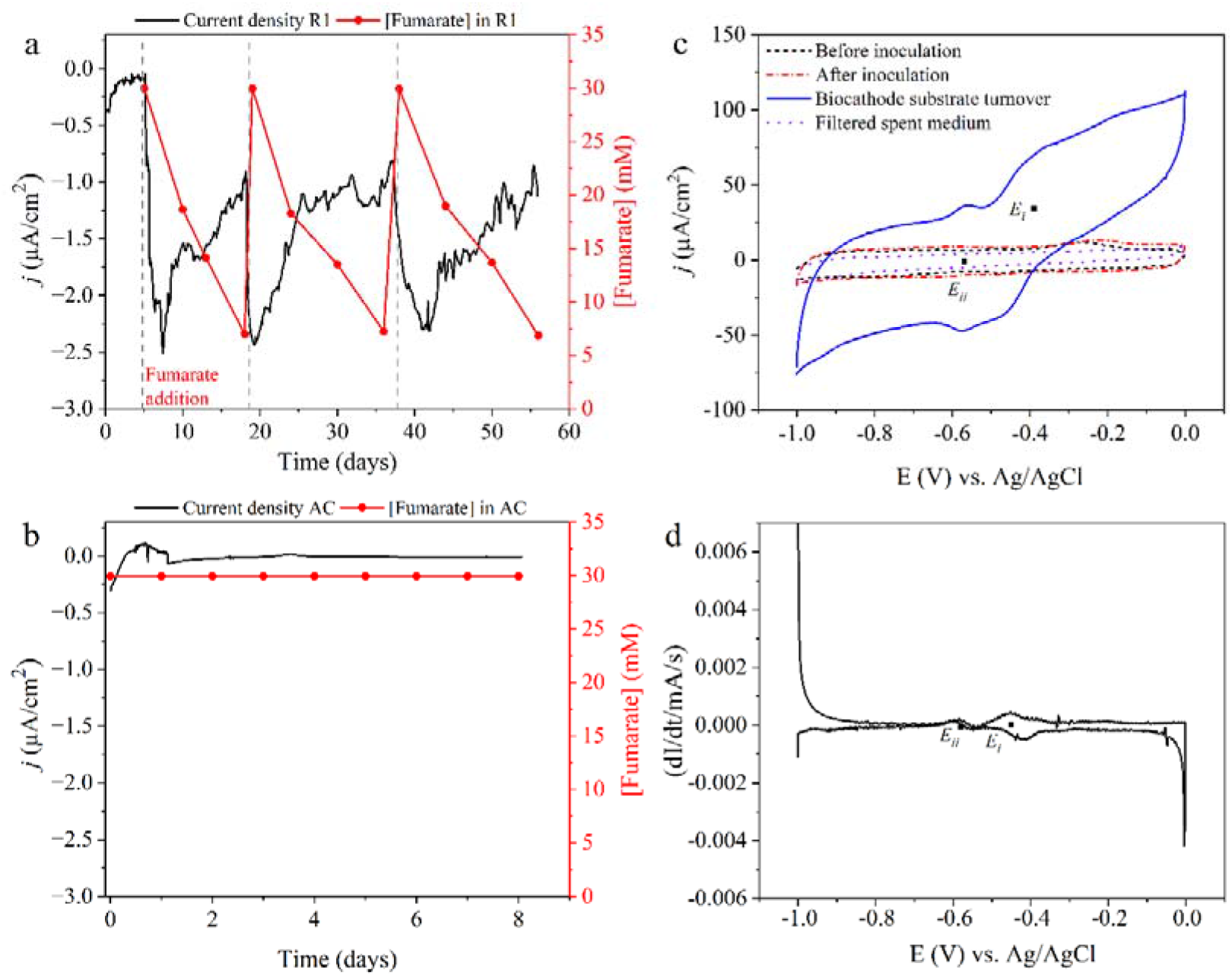
Electrochemical investigation of inward EET capability of *Geoalkalibacter halelectricus* with cathode as the sole electron donor (electron acceptor: fumarate). Bioelectrocatalytic cathodic current and fumarate concentration profiles of representative bioelectrochemical reactor R1 (a) and control experiments (b), cyclic voltammograms (CVs) at different conditions viz, before and after inoculation, during substrate turnover, and filtered cell-free spent medium (c), and the first derivative of CV recorded at substrate turnover condition (d). AC – Abiotic Control.

The SEM analysis of both the bioanode (**Figure S9a**) and biocathode (**Figure S9b**) confirmed the presence of a uniform and intact microbial biofilm comprising rod-shaped cells at the electrode surfaces. It supports the possibility of direct electron transfer and uptake by *G. halelectricus* biofilm from the anode and cathode, respectively.

## 4. Discussion

### 4.1 Bi-directional EET capability of *G. halelectricus*

Both biochemical and electrochemical cultivation approaches showed the outward and inward EET, and hence bi-directional EET capabilities of *G. halelectricus* under haloalkaline conditions. It is the first haloalkaliphilic electroactive microorganism known for the bi-directional EET. It uses a different set of redox moieties and, thus, different electron transfer components or processes in achieving outward and inward EET. Detailed characterization of the redox moieties involved in bi-directional EET processes would reveal the unique metabolic capabilities of this microbe. The high anodic current density and efficient Fe^3+^-oxide reduction reaction suggest the strong exoelectrogenic nature of *G. halelectricus*. In contrast, the low bioelectrocatalytic cathodic reduction current and limited and/or slow Fe^0^ oxidation reaction suggests the weak electrotrophic nature of *G. halelectricus*. It was able to draw only about −2.5±0.3 μA/cm^2^ reduction current, as reported previously for other electrotrophs, namely, *Geobacter sulfurreducens* (−3 μA/cm^2^) (Pous et al., 2016), *Acidithiobacillus ferrooxidans* (−0.5 μA/cm^2^) (Carbajosa et al., 2010), and *Morella thermoautotrophica* (−6.35 μA/cm^2^) (Yu et al., 2017). *Shewanella oneidensis* MR-1 has been previously reported to withdraw a maximum reduction current of −9 and −17 μA/cm^2^ from the cathode electrode via inward EET uptake mechanism with oxygen and fumarate (50 mM) as terminal electron acceptors, respectively (Ross et al., 2011; Rowe et al., 2018). *Geobacter sulfurreducens* have also been reported to withdraw a high current of −69 μA/cm^2^ for catalysing both energy-generating and storing reactions using fumarate as the terminal electron acceptor (Mickol et al., 2021). In this line, *G. halelectricus* needs to be tested with other electron acceptors such as nitrate, sulfate, and metal ions (Mn^3+^, Cr^6+^, Co^3+^, and Se^4+^) for its inward EET capabilities and to ascertain its electrophilic nature. It would eventually lead to a better understanding of its potential for microbial electrochemistry-based energy conversion and environmental remediation applications.

### 4.2 Iron cycling by *G. halelectricus* via bi-directional EET under haloalkaline conditions

The biochemical cultivation approach followed by the microscopic and EDS analysis of the *G. halelectricus* samples confirmed its capabilities to attach to the insoluble electron acceptor and donor surfaces to reduce Fe^3+^ oxide and oxidize Fe^0^ via outward and inward EET, respectively (Gupta et al., 2020; Kappler et al., 2021). These observations and the lack of any soluble redox mediator in the filtered cell-free spent medium suggested the direct mode of EET by *G. halelectricus* to transfer and uptake electrons in Fe^3+^ reduction and Fe^0^ oxidation reactions, respectively. These results suggest the involvement of *G. halelectricus* in iron cycling under haloalkaline conditions (Kato, 2016). Thus, its role can be implicated in increasing the bioavailability of Fe to other microorganisms or life forms under challenging extreme conditions and needs to be further investigated with different types of iron-bearing minerals (He et al., 2018; Nixon et al., 2022). *G. halelectricus* becomes the fourth haloalkaliphilic Fe^3+^-reducing microorganism along with *Fuchsiella ferrireducens, Bacillus arsenicoselenatis, and Bacillus selenitireducens* reported to date (Kappler et al., 2021; Nixon et al., 2022). Comparison of its Fe^3+^ reduction rate (i.e., 2.3 mM/day) with rates of *Fuchsiella ferrireducens* (1.7 mM/day) (Zhilina et al., 2015), the most studied haloalkaliphilic iron-reducer, suggests the crucial role of *G. halelectricus* in Fe cycling in haloalkaline environments. While the other haloalkaliphilic iron-reducers, namely *Bacillus arsenicoselenatis* and *Bacillus selenitireducens* have been quantitatively tested for Fe^3+^ reduction, but no reduction rates have been reported (Blum et al., 1998). The well-known iron-reducer, *Acidithiobacillus ferrooxidans* reduces iron at 6.7 mM/day rate under acidic conditions of pH 2 (Bhaskar et al. 2021). Notably, *G. halelectricus* iron reduction activity is on par with the other iron-reducers like *Geothrix fermentans* (1.9 mM/day) (Nevin and Lovley, 2002a), *Geobacter metallireducens* (1 mM/day), and *Shewanella alga* strain BrY (2.13 mM/day) (Nevin and Lovley, 2002b). These observations open up the opportunity to assess its performance in the anaerobic iron bioleaching process. Importantly, *G. halelectricus* is the first haloalkaliphile capable of both iron oxidation and reduction, and no other haloalkaliphile has been reported for both iron oxidation and reduction capabilities. **Figure 6** shows the putative role of *G. halelectricus* in iron cycling under haloalkaline conditions. The inward and outward EET capabilities suggest the occurrence of different metabolisms in *G. halelectricus*, i.e., the capability to switch on the respiration to insoluble electron donors and acceptors in environments lacking the readily available soluble electron donors and acceptors. Further investigations to understand the role of *G. halelectricus* and its genes involved in complete iron cycling via a direct mode of EET, besides its capabilities to respire on Fe^2+^-oxides as the sole electron acceptor or donor, are warranted.

**Figure 6:**
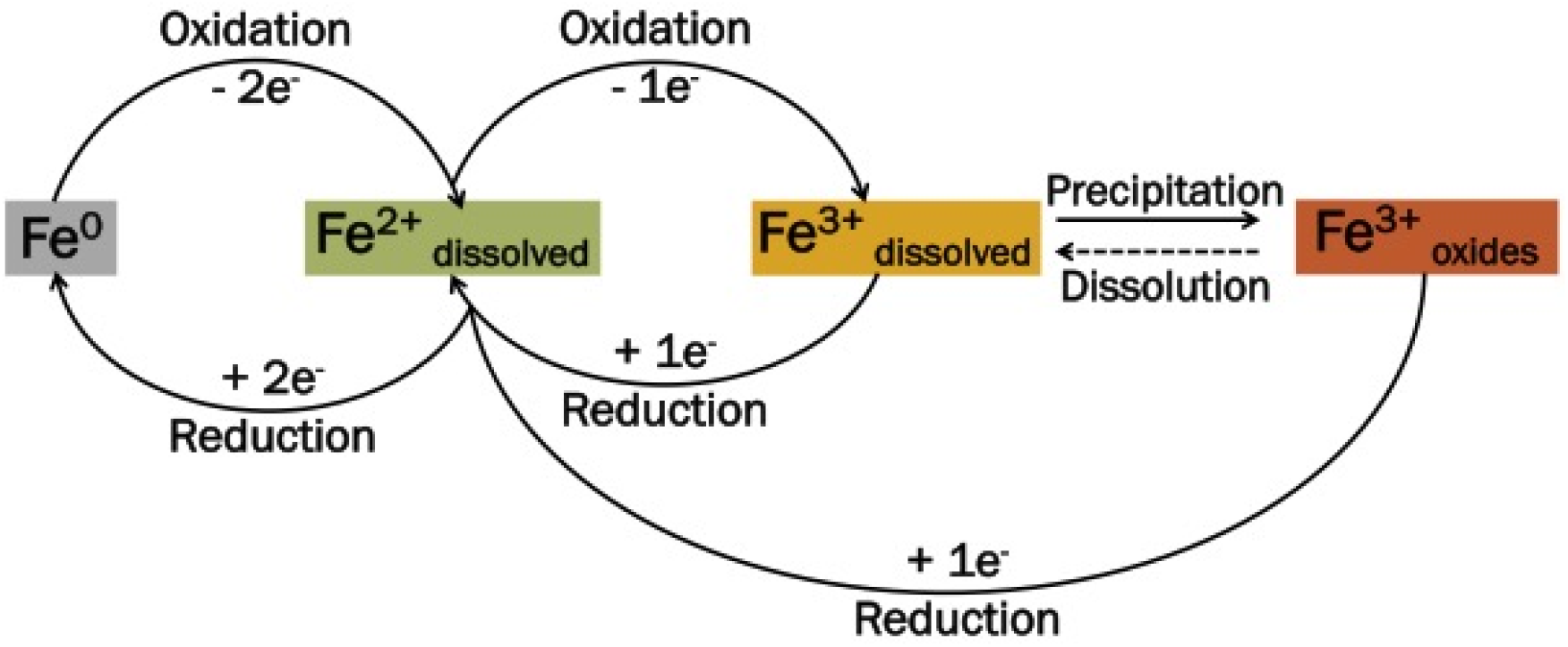
Schematic showing possible Fe cycling by *Geoalkalibacter halelectricus* under haloalkaline conditions.

## Conclusions

The serum-flask biochemical experiments revealed the capabilities of *G. halelectricus* to catalyse both Fe^3+^-oxide reduction and Fe^0^ oxidation via outward and inward EET, respectively, thereby, its bi-directional EET capabilities under haloalkaline conditions. The electrode-based bioelectrochemical cultivation further confirmed its capabilities to transfer electrons to the anode and draw electrons from the cathode via bi-directional EET. So far, no other extreme haloalkaliphilic microorganism has been reported for both iron oxidation and reduction capabilities. The electrochemical investigation of the *G. halelectricus* biofilms and filtered cell-free spent media by cyclic voltammetry revealed the involvement of redox-active components with different mid-point potentials and lack of soluble redox mediators in outward and inward EET processes. This study on a haloalkaliphilic microorganism capable of bi-directional EET contributes to the limited understanding of this important microbial EET phenomenon. It also opens up opportunities to understand the relevance of bi-directional EET to key natural processes, such as iron cycling in extreme environments.

## Acknowledgements

Sunil A. Patil is grateful to IISER Mohali for the infrastructural and financial support. Sukrampal Yadav acknowledges IISER Mohali for the Ph.D. fellowship. Chetan Sadhotra (IF190662) acknowledges the Department of Science and Technology-Innovation in Science Pursuit for Inspired Research (DST-INSPIRE), Government of India, for the doctoral fellowship.

